# Reducing SAR in 7T brain fMRI by circumventing fat suppression while removing the lipid signal through a parallel acquisition approach

**DOI:** 10.1101/2020.10.13.337691

**Authors:** Amir Seginer, Edna Furman-Haran, Ilan Goldberg, Rita Schmidt

## Abstract

Ultra-high-field functional magnetic resonance imaging (fMRI) offers the way to new insights while increasing the spatial and temporal resolution. However, a crucial concern in 7T human MRI is the increase in power deposition, supervised through the specific absorption rate (SAR). The SAR limitation can restrict the brain coverage or the minimal repetition time of fMRI experiments. fMRI is based on the well-known gradient-echo echo-planar imaging (GRE-EPI) sequence, which offers ultrafast acquisition. Commonly, the GRE-EPI sequence comprises two pulses: fat suppression and excitation. This work provides the means for a significant reduction in the SAR by circumventing the fat-suppression pulse. Without this fat-suppression, however, lipid signal can result in artifacts due to the chemical shift between the lipid and water signals. Our approach exploits a reconstruction similar to the simultaneous-multi-slice (SMS) method to separate the lipid and water images, thus avoiding undesired lipid artifacts in brain images. The lipid-water separation is based on the known spatial shift of the lipid signal, which can be detected by the multi-channel coils sensitivity profiles. Our study shows robust human imaging, offering greater flexibility to reduce the SAR, shorten the repetition time or increase the volume coverage with substantial benefit for brain functional studies.

## Introduction

Ultra-high field (≥7T) magnetic resonance imaging (MRI) offers both an increased signal-to-noise ratio (SNR) and an improved contrast-to-noise ratio (CNR), which can be exploited to increase the spatial and temporal resolution. One of the methods that has harnessed these benefits is functional MRI (fMRI). fMRI is based on a well-known gradient-echo echo-planar imaging (GRE-EPI) sequence, which offers ultrafast acquisition^1^. However, EPI is also noted for its low effective bandwidth in the phase encoding (PE) direction^2^. One of the consequences of this low bandwidth is the introduction of a large lipid-water shift in the image due to the chemical shift between the lipid and water signals, resulting in artifacts in the image. The shift proportionally increases with the strength of the magnetic field. The lipid artifacts can reduce image quality and fMRI efficiency, particularly since fMRI is based on the detection of small signal changes and is therefore sensitive to undesirable artifacts. An effective technique commonly used to remove these artifacts is to prepend an RF pulse to suppress the lipid signal. Therefore, a common GRE-EPI sequence is comprised of two pulses - fat suppression and excitation. When targeting whole-brain coverage (with a high slice resolution) and a reduced repetition time, fMRI in 7T can reach the limits of the allowed specific absorption rate (SAR). The fat-suppression pulse can double and even triple the total SAR, depending on the scan parameters. In addition, increased RF field inhomogeneity in ultra-high field MRI can locally reduce the efficiency of fat suppression.

Several works have sought to improve the fat-suppression techniques by addressing the RF field inhomogeneity and by reducing the SAR. These studies include SAR-optimized pulses and adiabatic pulse implementations ^3,4,5,6^. A wide range of other efforts have been directed to developing techniques for the removal of the lipid signal, such as inversion recovery pulses^7^, water excitation^8^ and opposite sign gradient in spin-echo implementation^9,10^. Recent studies have explored the use of methods based on fingerprinting^11,12^ and compressed sensing^13,14^ to reconstruct separate images of the lipid and water.

In this study, we examine the potential to significantly reduce the SAR by circumventing the fat-suppression pulse. To complement the removal of the fat-suppression pulse, we utilized a reconstruction based on the parallel acquisition technique to separate the lipid and water images. An EPI implementation without fat suppression can offer fMRI studies greater flexibility to reduce the SAR, shorten the repetition time or increase the volume coverage.

Since the introduction of SENSE^15^ and GRAPPA^16^, parallel acquisition methods that accelerate the scan and shorten the echo time have proven highly efficient. Reconstruction methods are constantly being improved, including implementations such as ESPIRIT^17^ and simultaneous multi-slice for EPI with CAIPIRINHA^18^. Moreover, the efficiency of parallel acquisition is further enhanced by advances in coil technology and increase in the magnetic field^19^. SENSE and GRAPPA based techniques have also been examined in applications other than acceleration. Examples include the correction of ghost artifacts in EPI^20^ and restricted FOV imaging^21,22^. Parallel acquisition has also been shown to improve the reconstruction in chemical shift imaging^23^ and in lipid ghosts elimination^24^. The SENSE method has been successfully applied in hyperpolarized ^13^C metabolic imaging to reconstruct separate metabolite images^25^; based on their chemical shift. In several feasibility studies SENSE has also been demonstrated to separate lipid and water images from EPI acquisitions at 1.5 T and 3 T MRI^26–28^. In the current study, we further explore lipid water separation for EPI in 7T MRI. The separation is based on the distinct chemical/spatial shift of the lipid signal that can be detected by the multi-channel coils’ sensitivity profiles. This can be visualized as analogous to CAIPIRINHA’s^29^ shift of the slices in simultaneous-multi-slice (SMS) acquisition. In the lipid-water case, the introduced FOV shift is defined not by the added gradient blips but rather by the bandwidth in the PE direction and the known chemical shift of the lipid peak (3.4 ppm which are ~1000 Hz at 7T). The arising large lipid-water signal shift in the EPI images actually aids reliable reconstruction.

In this study, lipid-water separation is examined in combination with parallel acceleration – including both in-plane and SMS. The benefits of the proposed strategy are examined, including the substantial reduction in the SAR. One of the concerns is the potential impact of the lipid-water separation on the accessible acceleration factor. To examine this issue, we estimated the geometry factor (g-factor) maps for lipid-water separation compared to SMS acceleration. In addition, human volunteer scanning was performed to examine the reconstruction quality in-vivo and to assess fMRI efficiency. Temporal SNR (tSNR) was compared in experiments with and without fat-suppression pulse.

### Lipid and water images separation: Principles and extended parallel imaging formulation

Lipid-water images can be reconstructed separately using the parallel imaging technique if distinct sensitivity profiles exist as was demonstrated in Ref.^28^. The combined lipid-water acquired signal can be described as follows:

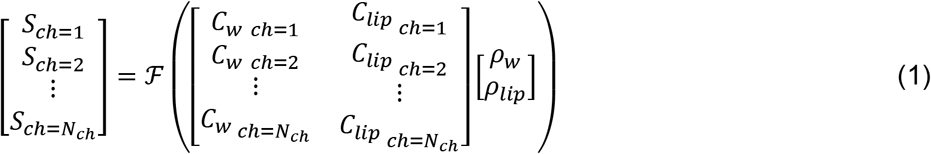

where *S*_*ch*_ is the k-space acquired signal per channel (*ch*=*1*‥*N*_*ch*_), 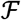 is the FFT operator, *C*_*lip*_ and *C*_*w*_ are the per-channel sensitivity profiles of the lipid and water signals, respectively, and *ρ*_*lip and*_ *ρ*_*w*_, are the lipid and water images, respectively. *C*_*lip*_ can be generated by either shifting the *C*_*w*_ by the known lipid-water spatial shift or by acquiring an additional lipid image (e.g., by using a water suppression pulse) and shifting it by the same known lipid-water shift. Note that this formulation is analogous to SMS reconstruction.

The shift of lipid (*d*_*lip-w*_) in EPI is defined as follows:

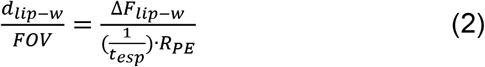

where *FOV* – is the FOV along the PE direction, Δ*F*_*lip−w*_ is the spectral shift of lipid from water in Hz (3.4 ppm, which are ~1000Hz at 7T), *t*_*esp*_ is the echo spacing in seconds and *R*_*PE*_ is the acceleration factor along the PE direction, if applied.

Table 1 shows the spatial shifts for representative sets of EPI scans. A higher in-plane acceleration rate proportionally reduces the shift. Notice that the shifts for commonly used scan parameters are substantial, which, as shown here, can be exploited in favor of a reliable separate reconstruction of lipid and water images.

**Table 1:**
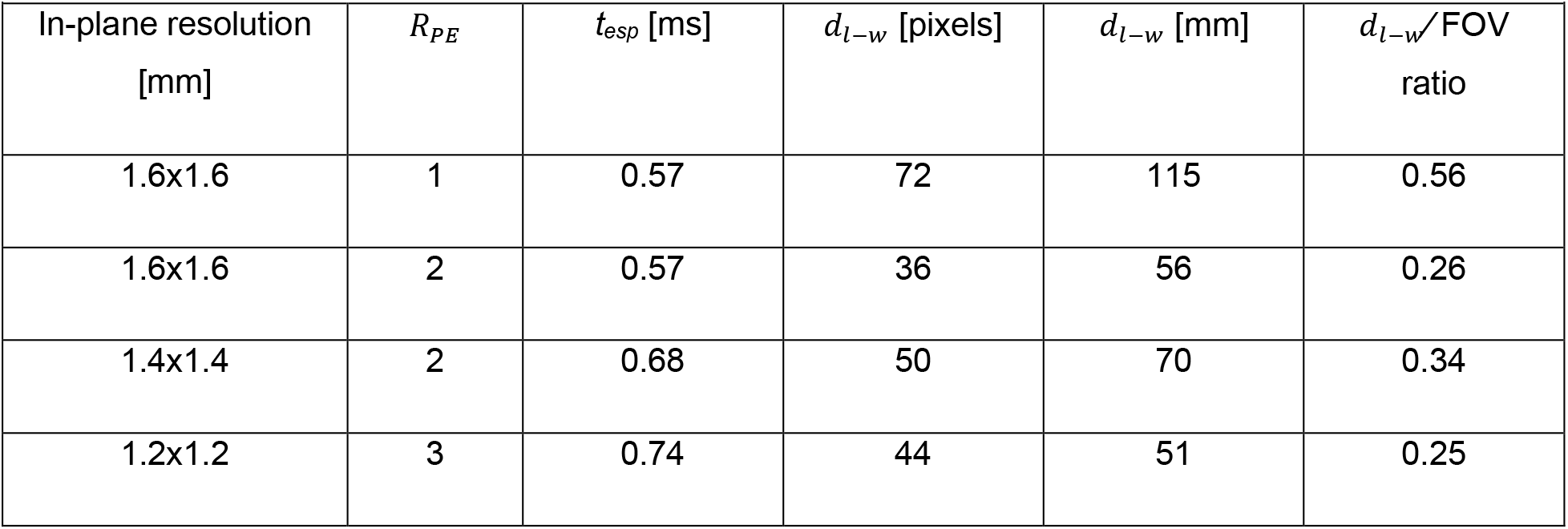
Lipid-water shift for representative EPI parameters.

| **Table 1: Lipid-water shift for representative EPI parameters** |  |  |  |  |  |
| --- | --- | --- | --- | --- | --- |
| In-plane resolution<br>[mm] | $R_{PE}$ | $t_{esp}$ [ms] | $d_{l-w}$ [pixels] | $d_{l-w}$ [mm] | $d_{l-w}/FOV$<br>ratio |
| 1.6x1.6 | 1 | 0.57 | 72 | 115 | 0.56 |
| 1.6x1.6 | 2 | 0.57 | 36 | 56 | 0.26 |
| 1.4x1.4 | 2 | 0.68 | 50 | 70 | 0.34 |
| 1.2x1.2 | 3 | 0.74 | 44 | 51 | 0.25 |

To implement a “fit for all” reconstruction method for all three parallel imaging aspects – (i) in-plane PE acceleration, (ii) SMS acceleration, and (iii) lipid-water separation - the problem was reformulated in a manner that treats all three aspects as if they were case (i), i.e., in-plane PE acceleration. This is achieved by solving a “full-FOV” image defined as the concatenation of all final images (all the slices of the separate water and lipid images), similarly to previous works without lipid-water separation^30,31,32^. Notice that this extended formulation, further detailed below, was established for convenience and supports any of the developed and well-established reconstruction methods for parallel imaging. This formulation simplifies the reconstruction procedure, since it utilizes the same engine to solve the inverse problem. The prerequisites for reconstruction are only the input signal and a set of sensitivity maps.

For simplicity, let us first describe the case of the acquisition of *R*_*sms*_ simultaneous slices together with an in-plane PE acceleration. The acquired signal per channel can be described in the following manner:

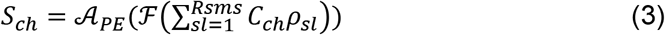

where 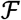 is the FFT operator, 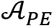 is a PE subsampling operator mimicking the actual acquisition that samples only every *R*_*PE*_ line, and 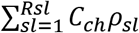 is the sum of the simultaneously acquired slices, where *sl*=*1*‥*R*_*sms*_ is the slice counter. In a matrix form, the dimension of *S*_*ch*_ is *(N*_*PE*_∕*R*_*PE*_) x *N*_*RO*_, where *N*_*PE*_ and *N*_*RO*_ are the number of points along the PE and RO directions, respectively.

In the extended formulation proposed here, we first stack the slices in the PE dimension, to form an *(R*_*sms*_·*N*_*PE*_) x *N*_*RO*_ matrix and then the final signal is subsampled by an acceleration factor *(R*_*sms*_·*R*_*PE*_), represented by operator 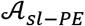. This is represented by the following equation:

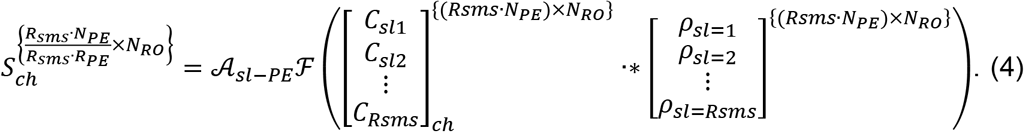

The matrix sizes are indicated here in the right upper corner. The.* stands for element-by-element multiplication and *C*_*sl*_ are per-channel sensitivity profiles of the slices.

Having acquired the signal (left-hand side) and the sensitivity maps, we can solve the inverse problem of Eq. 4 to estimate the slice images (*ρ*_*sl*=1,_*ρ*_*sl*=2_, ‥, *ρ*_*sl*_=*R*_*sl*_), just like in a regular in-plane parallel imaging problem. Note that this holds for an odd number of slices. For even number of slices, the whole stack of slices has to be shifted by a half-FOV, to center a slice at the center of the new effective (stacked) image. This is explained schematically in Figure 1 and in more detail in the Supporting Information. Therefore, for an even number of slices, concatenating the slices will give us:

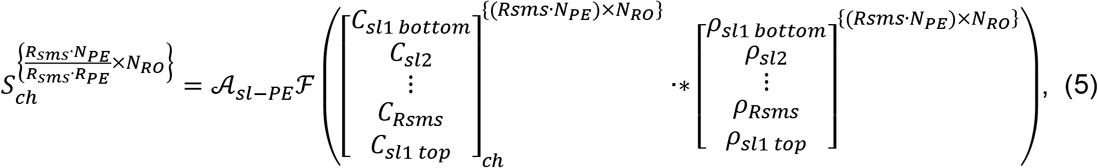

where “bottom” and “top” refer to the bottom and top halves of the image. Notice that in this formulation, the simultaneous acquisition of two slices is analogous to an in-plane PE acceleration by a factor of two. The description above does not include CAIPIRINHA implementation. If CAIPIRINHA is used, the CAIPIRINHA shift is applied separately for each slice and its sensitivity map to match the applied shift during the scan.

**Figure 1:**
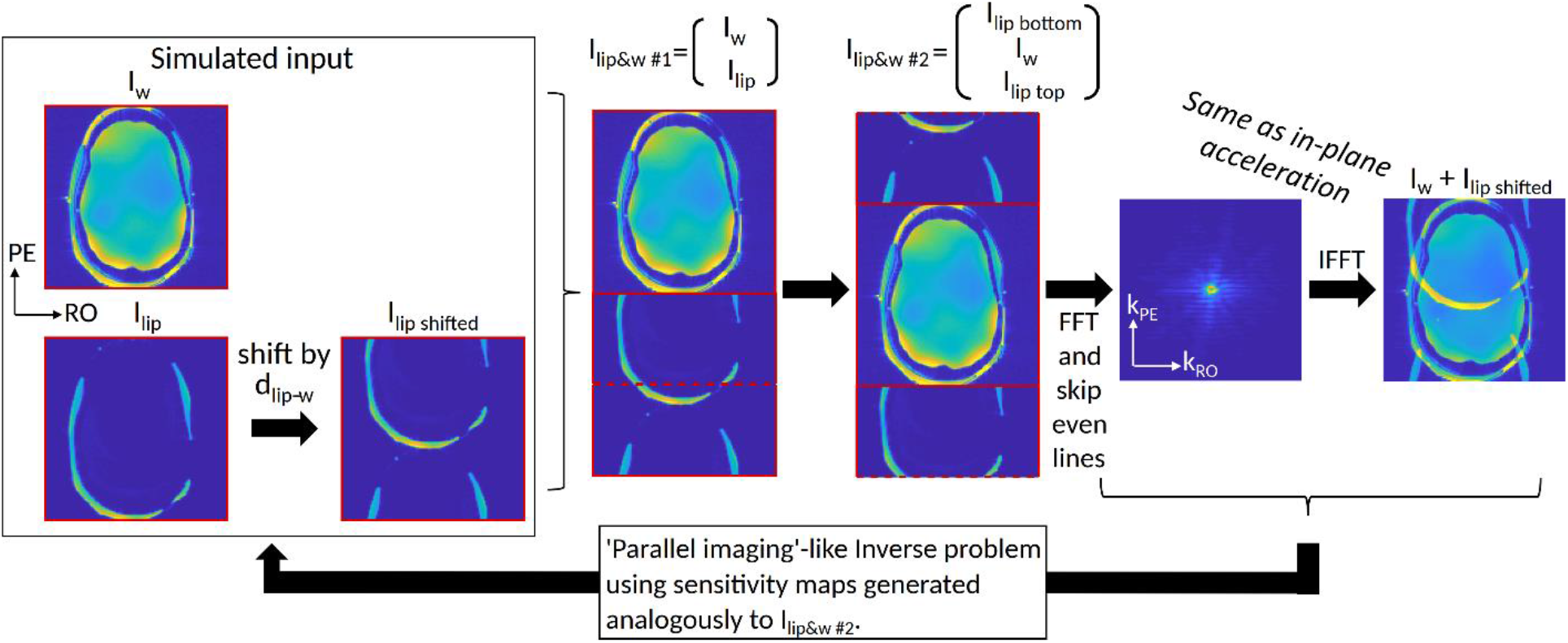
Extended formulation that can combine three parallel imaging aspects (i) in-plane PE acceleration, (ii) SMS acceleration and (iii) lipid-water separation. The steps show how a resulting EPI image can be described for water and lipid “slices” (I_w_ and I_lip_). The I_w_ and I_lip_ shifted are concatenated, then shifted together by a half-FOV. Images “I_w_+I_lip shifted_” (at the far right) and “I_w_&_lip #2_” resemble the case of in-plane acceleration, with “I_w_+I_lip shifted_” analogous to the acquired image (due to sub-sampling) and “I_w_&_lip #2_”analogous to the full-FOV image in such a case. Following this description, one can solve the inverse problem analogous to in-plane parallel imaging using sensitivity maps. The same description holds also for two slices acquired simultaneously (instead of lipids and water signals). Finally, the formulation can be extended to any number of slices and fat/water images.

Analogously to Eq.3 for SMS, the lipid-water separation signal can be written as:

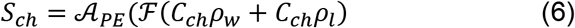

and therefore, in analogy to Eq. 4, can also be described as:

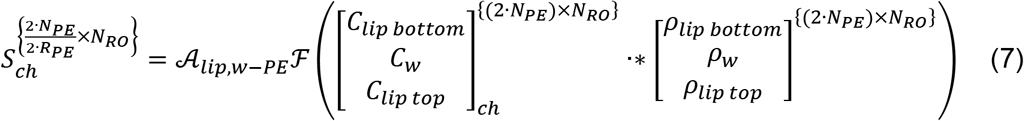

where 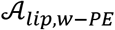 is the subsampling operator that mimics the actual acquisition, sampling both the lipid and the water images and, therefore, subsamples every 2·*R*_*PE*_ lines (2 for lipid and water). *C*_*ch*_*ρ*_*w*_ + *C*_*ch*_*ρ*_*l*_ is the sum of the combined lipid and water signal.

Finally, the lipid-water and slices datasets can be jointly concatenated to give a total effective “acceleration” factor of *R*_*tot*_=*2R*_*sms*_·*R*_*PE*_. For example, the case of *R*_*sms*_=2 with lipid-water separation will be analogous to acceleration by a factor of four, *R*_*tot*_=2·2·1.

With this formulation in hand, the solution can be based on any parallel imaging reconstruction method, including the well-known SENSE, GRAPPA, ESPIRIT, as well as any other inverse problem solution. In this study, a custom-written MATLAB code was used to arrange the input k-space and the sensitivity profiles. The final reconstruction was performed utilizing the BART^33^ software, using the “pics” command with L1 norm.

## Results

Phantom experiments were conducted to examine the reconstruction quality. A 3D-printed head phantom that included lipid and brain compartments was used. Figure 2 compares the lipid-water separation with R_sms_=2. The same slice is reconstructed in both cases and the g-factor maps for this slice are compared. The average and standard deviation of the g-factor are 1.06 0.12 for lipid-water separation and 1.01±0.08 for R_sms_=2. The maximum g-factors inside the phantom are 1.6 and 1.4 for lipid-water separation and R_sms_=2, respectively. The low g-factor for R_sms_=2 is due to the CAIPRINHA implementation. Unlike the SMS case, the lipid-water g-factor map shows a localized increase in a narrow range, that of the shifted lipid layer region. The SNR of the image without fat suppression was estimated to be x1.3 higher than with fat suppression.

**Figure 2:**
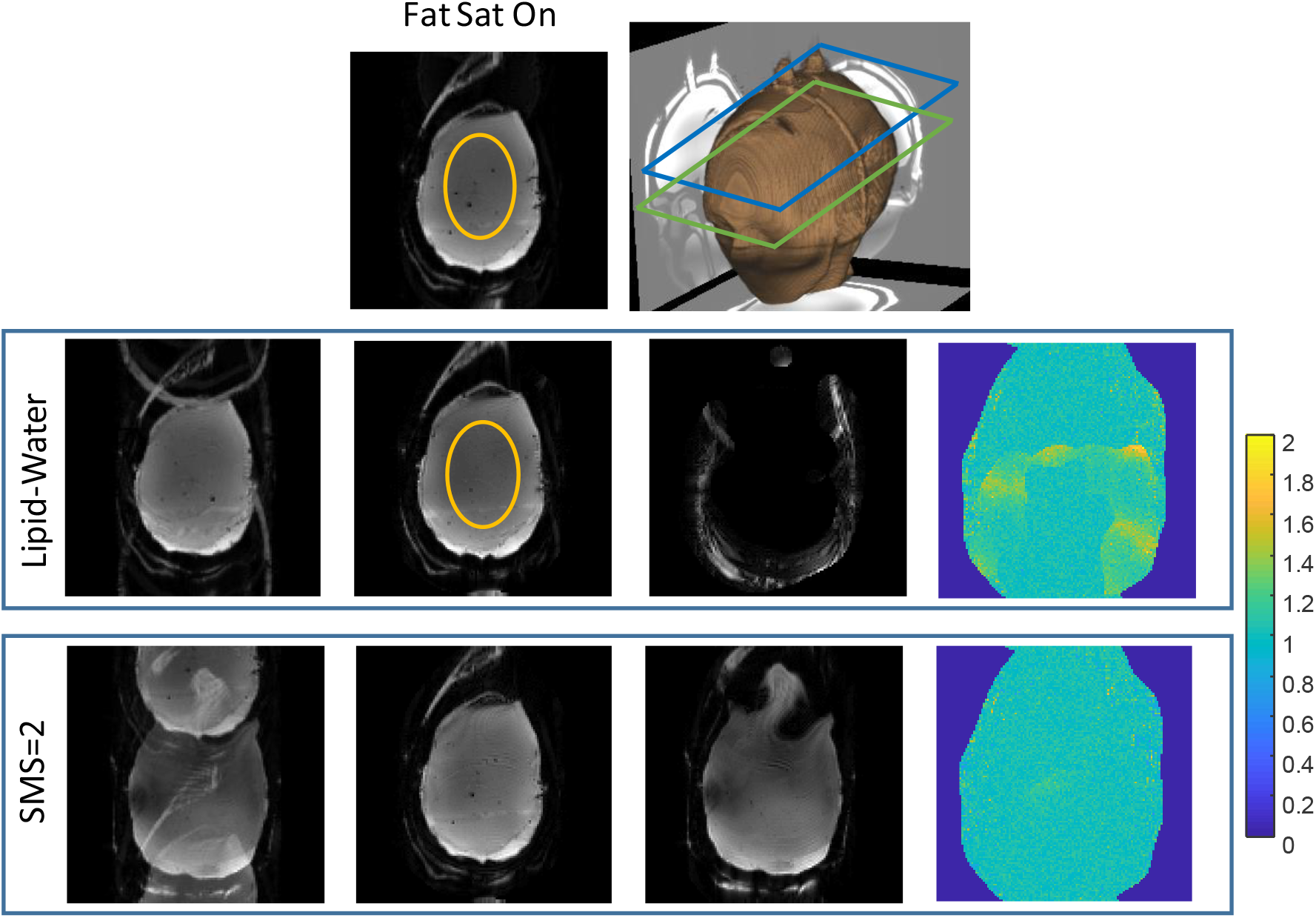
Phantom scanning – comparison of Lipid-Water and SMS reconstruction. Top row - Image acquired with fat suppression and a 3D rendering showing the phantom shape, the main slice location (green frame) and a second slice (blue frame). Second row - Images acquired without fat suppression. From left to right – image with standard reconstruction (with overlapping lipid signal), reconstruction of separated water and lipid images, and a g-factor map for the water image. Bottom row - Images acquired with R_sms_=2. From left to right – image with standard reconstruction (with overlapping slices), reconstructed two slices images and g-factor map for the first slice. The ROIs shown in the orange overlay were used to estimate the SNR. The g-factor color-map range is 0-2.

Figure 3 shows a further comparison of combined acceleration factors with lipid-water separation. It includes well-reconstructed images with a total acceleration of *R*_*tot*_=2 (lipid-water separation only), *R*_*tot*_=4 (lipid-water separation and *R*_*PE*_ =2) and *R*_*tot*_=8(lipid-water separation, *R*_*PE*_ =2 and R_sms_=2). The g-factor values increase to 1.18±0.26 for *R*_*tot*_=4 and to 1.31±0.28 for *R*_*tot*_=8.

**Figure 3:**
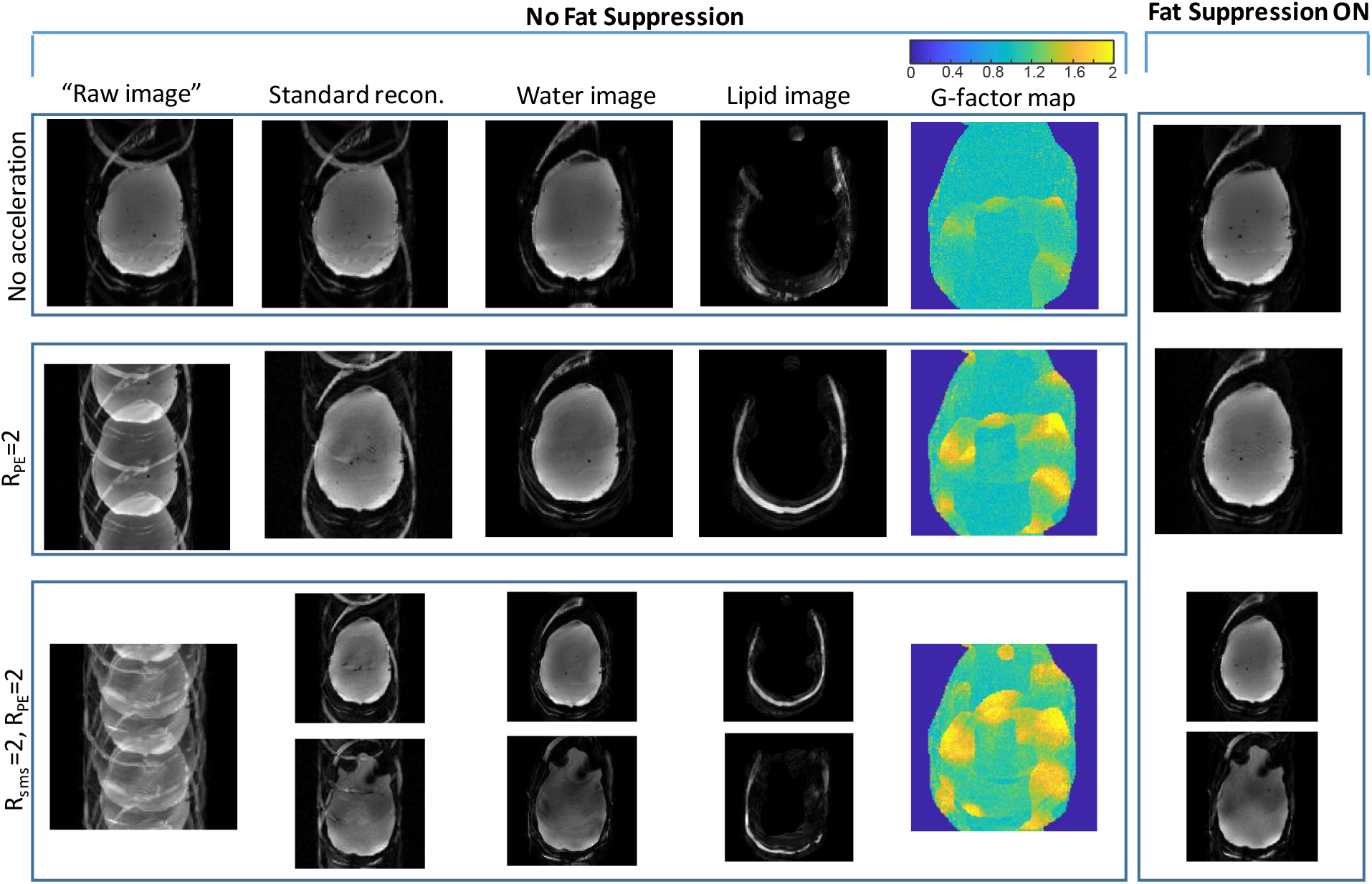
Phantom scanning – combining in-plane acceleration, SMS and lipid-water separation. From top to bottom - *R*_*tot*_=2 no acceleration (Lipid/Water separation), *R*_*tot*_=4 (Lipid/Water, *R*_*PE*_ =2), *R*_*tot*_=8 (Lipid/Water, *R*_*PE*_ =2, R_sms_=2). From left to right – “raw image” (FFT applied directly to the acquired image), “standard recon.” (Siemens product reconstruction), images reconstructed with the extended formulation - separate water and lipid images, g-factor maps for the common slice, and the image acquired with fat suppression. The g-factor color-map range is 0-2.

Human volunteers were scanned to examine the lipid-water separation in-vivo and to verify the fMRI efficiency. Figure 4 and 5 demonstrate human imaging of two representative slices out of 30 well-reconstructed slices for scans with and without fat-suppression. Figure 4 shows two scans with in-plane resolution of (a) 1.6 mm and (b) 1.4 mm. As increasing the resolution commonly requires higher in-plane acceleration, we examined *R*_*PE*_ =2 for the first case (1.6 mm) and *R*_*PE*_ =3 for the second case (1.4 mm). The lipid signal artifact can be observed in the standard reconstruction without fat suppression (see yellow arrows). Examples of a relative signal profile (relative to the fat-suppression case) along two lines are also shown (denoted “1” and “2”). The lipid artifact reaches 35% along line “1”and 30% along line “2” and is practically removed after lipid separation. An additional experiment combining both in-plane and slice acceleration – *R*_*PE*_ =2 with either R_sms_=3 or R_sms_=4 – is summarized in Figure 5. The lipid signal artifact, in these cases, can be observed in the standard reconstruction (see yellow arrows). Water images with R_sms_=3 (R_tot_=12) are well-reconstructed, while R_sms_=4 (R_tot_=16) introduce some artifacts (see blue arrows).

**Figure 4:**
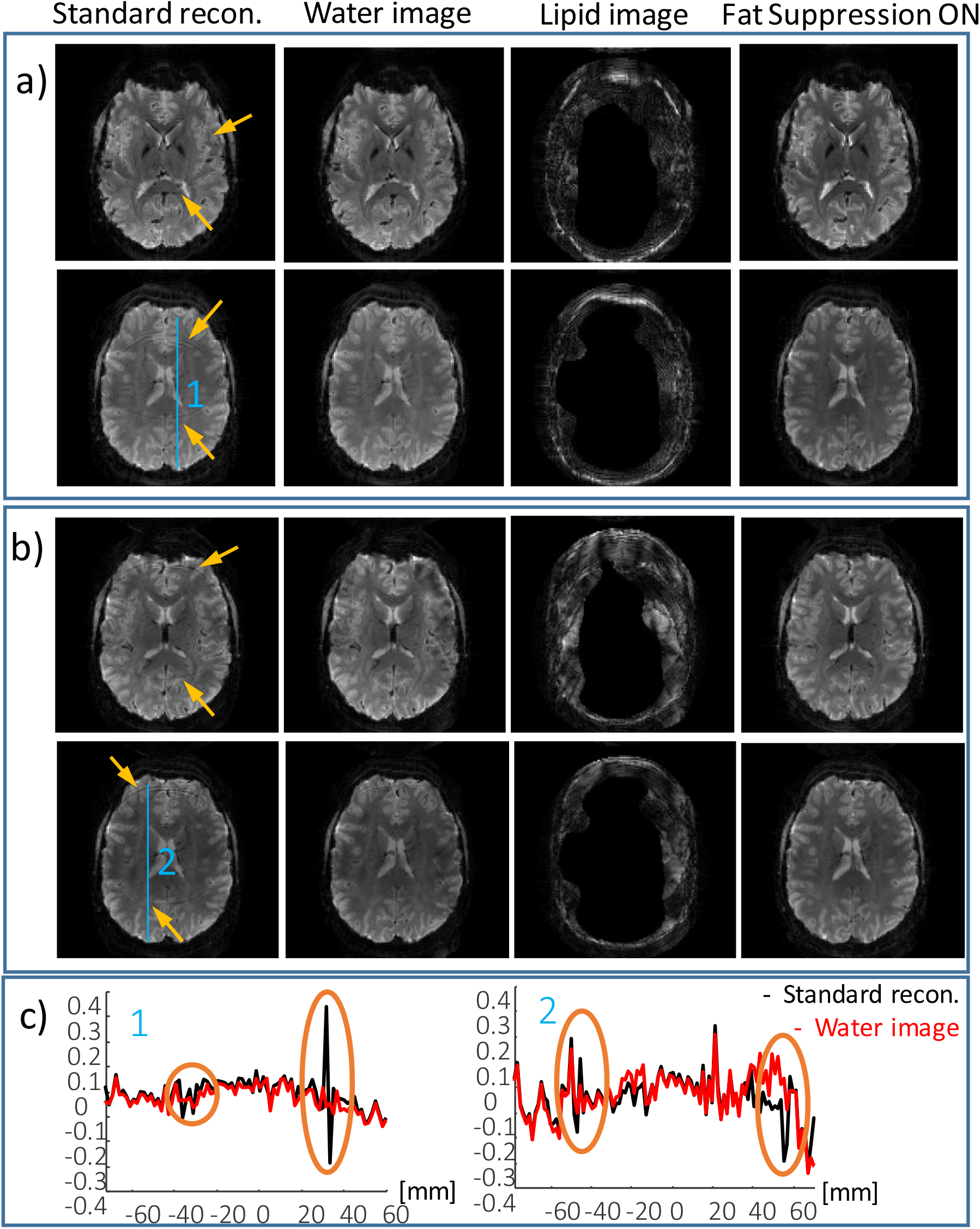
Human imaging – combined in-plane acceleration and lipid-water reconstruction. Two representative slices are shown for *R*_*PE*_ =2 (a) and *R*_*PE*_ =3 (b). From left to right -standard recon. (Siemens product reconstruction), water image, and lipid image from scan without fat suppression, and the image acquired with fat suppression. (c) Relative signal change 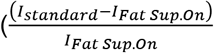 and 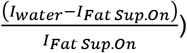 along the blue lines 1) and 2) shown in (a) and (b), respectively. Yellow arrows point to the artifact due to the lipid signal in the standard reconstruction.

**Figure 5:**
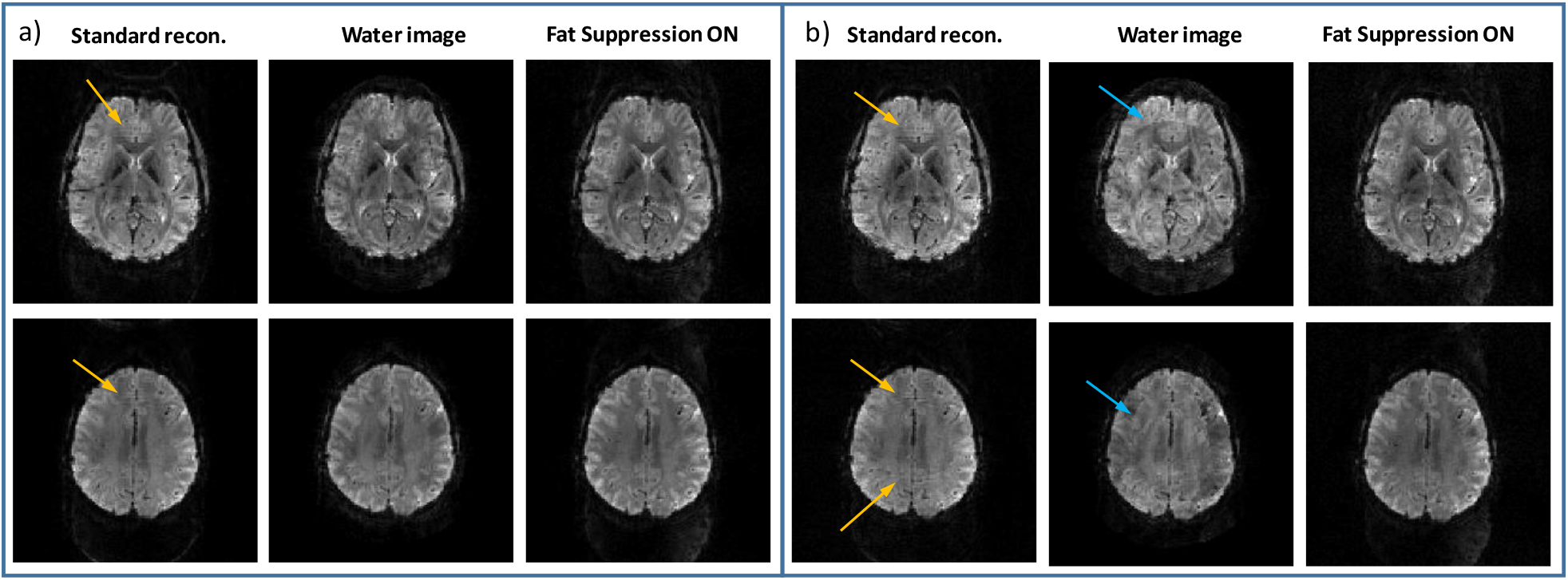
Human imaging – combined in-plane acceleration, SMS and lipid-water reconstruction. Two representative slices are shown for (a) *R*_*PE*_ =2 and R_sms_=3 and (b) *R*_*PE*_ =2 and R_sms_=4. From left to right – standard recon. (Siemens product reconstruction), water image, and lipid image from scan without fat suppression, and the image acquired with fat suppression. Yellow arrows point to the artifact due to the lipid signal in the standard reconstruction. Blue arrows point to artifacts that occur due to high acceleration factor - R_tot_.

In addition, scan parameters for fMRI experiments were examined to identify SAR restrictive cases. Table 2 shows a comparison of SAR levels between fMRI experiments with and without fat suppression. At 7T, fMRI experiments commonly require in-plane acceleration of factor 2 or 3 to achieve optimal TE (in the range of 20-35 ms, depending on the region of interest). We examined parameters for scans with resolution and slice coverage of interest for fMRI, targeting TE in the above range and with TR in the range of 1.5-2 seconds, a typically desired temporal resolution. Although SAR is reduced when increasing the SMS factor, in practice, combining in-plane acceleration with higher SMS factors introduces artifacts and reduces SNR – depending on the specific receive coil – and therefore SMS factors of 2 or 3 are of interest. Table 2 shows examples with SMS of factors 2 and 3, which are SAR constrained when fat suppression is applied, but are significantly relaxed if fat suppression is avoided. Cases 1-3 are based on the Siemens product EPI sequence and cases 4-5 on the multi-band GRE-EPI sequence from the University of Minnesota Center for Magnetic Resonance Research (CMRR) ^31,34^, which was examined since its RF excitation pulse can be shortened and so achieve a shorter TE (which is required for higher resolution scans).

**Table 2:** Examples of scan parameters with restrictive SAR. (the reference amplitude for 1 ms 180° hard pulse: 240 V)

|  |  |  |  |  |  |  |  |  |
| --- | --- | --- | --- | --- | --- | --- | --- | --- |
| Sequence | Case | $N_{SI}$ | $R_{PE}$ | $R_{SMS}$ | TE [ms] | TR [ms] | SAR [%]<br>Fat. Sup. ON | SAR [%]<br>Fat. Sup. OFF |
| Ep2d_bold<br>(Siemens product)<br>1.6 mm iso,<br>flip angle=90 | 1 | 64 | 3 | 2 | 20<br>(partial<br>Fourier<br>=7/8) | 1500 | 101 | 34 |
|  | 2 | 60 | 3 | 2 | 21 | 1500 | 97 | 33 |
|  | 3 | 76 | 3 | 2 | 21 | 1500 | * | 35 |
| cmrr_mbep2d_bold<br>1.4 mm iso,<br>RF excitation<br>pulse dur. = 4 ms,<br>flip angle 90 | 4 | 78 | 3 | 2 | 31.6 | 2000 | 122 | 80 |
|  | 5 | 78 | 3 | 3 | 31.6 | 2000 | 101 | 57 |
| * This case could not be acquired with TR shorter than 1860 ms (because of the extra time required for the fat suppression pulses)' |  |  |  |  |  |  |  |  |

The human scans also included a five minute-long (200 repetitions) resting state fMRI, both with and without fat suppression, to estimate the SNR and tSNR for each case (summarized in Figure 6). In this test, a combination of *R*_*PE*_ =3 and R_sms_=2 was applied to achieve a spatial and temporal resolution representative of a 7T-repetition time of 1.5 seconds and an isotropic resolution of 1.7 mm. This acceleration results in *R*_*tot*_=12 when lipid-water separation is included in the reconstruction. A pair of simultaneously acquired slices are shown without and with lipid-water separation compared to applying fat-suppression. In (c) tSNR maps without and with fat-suppression are compared. The SNR ratio of R_tot_=12 (*R*_*PE*_ =3, R_sms_=2, Lipid-Water) without fat suppression to R_tot_=6 (*R*_*PE*_ =3, R_sms_=2) with fat suppression, were 1.35 and 1.23 in regions 1 and 2, respectively. The tSNR ratio was 1.47 and 1.13 in regions 1 and 2, respectively.

**Figure 6:**
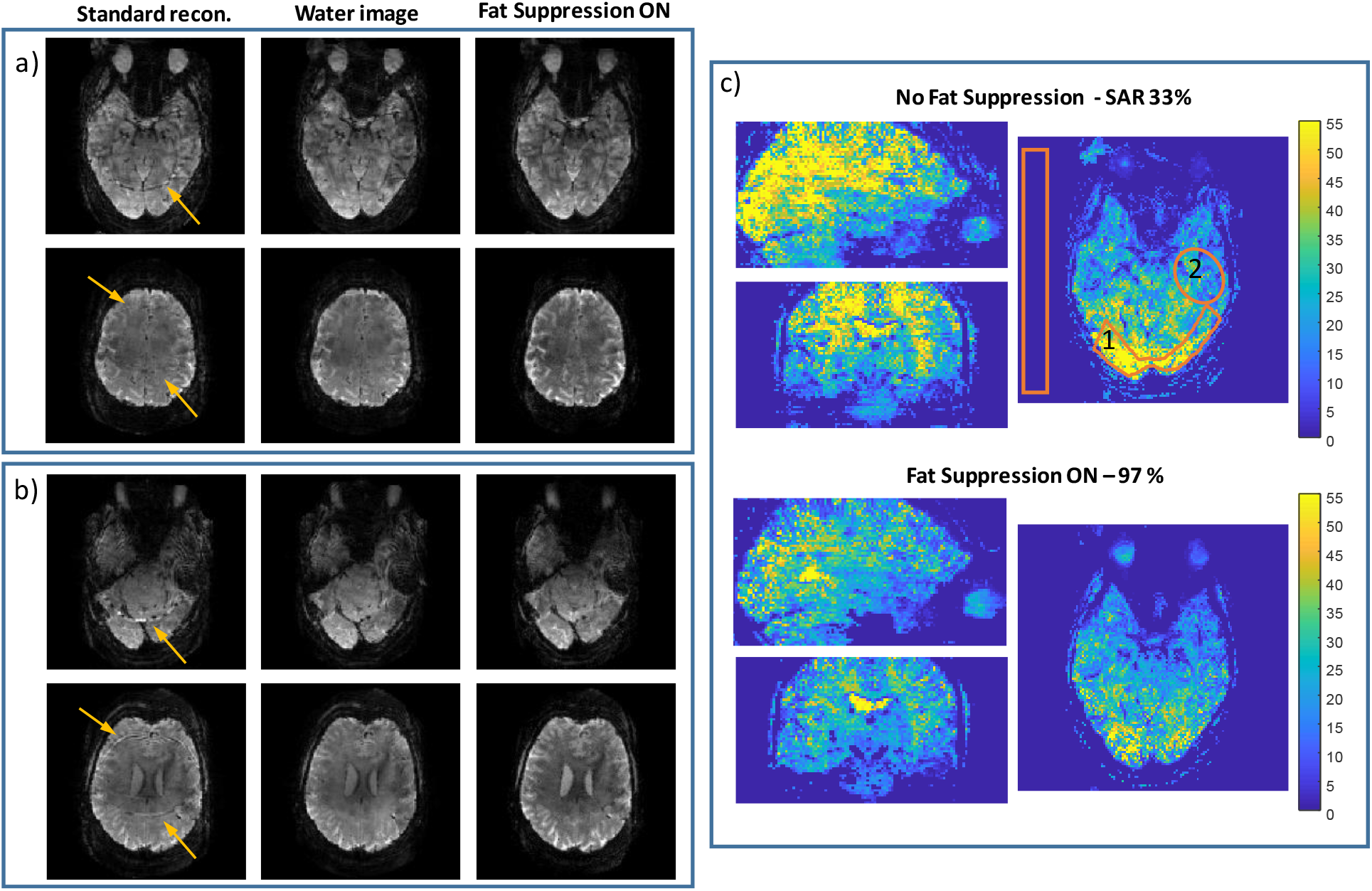
Human imaging – resting-state fMRI scan. Reconstruction of two pairs (a and b) of simultaneously acquired slices; comparing, left to right, standard reconstruction, a water image after lipid separation and an image acquired with fat suppression. (c) tSNR comparison in three orthogonal planes. Yellow arrows in (a) and (b) point to the artifact due to the lipid signal in the standard reconstruction. The orange overlays in (c) show regions (1) and (2) for the SNR and tSNR estimation. The SAR was 33% (without fat-suppression) and 97% (with fat-suppression) where the reference amplitude for 1 ms 180° hard pulse - 240 V.

## Discussion

Pushing the limits and moving to 7T MRI allows to increase the spatial resolution as well as to shorten the repetition time. However, scanning at 7T also increases the SAR, making it a limiting factor, including in fMRI^35^. Methods to reduce the SAR are, therefore, in high demand. In this study, we demonstrate a method that allows to avoid fat suppression in EPI, a method with which the SAR of a GRE-EPI can be reduced by a factor of two or even three. This implementation allows better flexibility in the design of the scan protocol: reducing the SAR, shortening the repetition time or increasing the slice coverage. fMRI experiments commonly require long repeating scans with a total duration above 30 minutes. They can therefore benefit tremendously from reducing the total SAR of the experiment. In addition, Table 2 shows examples in which circumventing fat suppression will achieve a larger number of slices or shorter TR, with resolution and slice coverage that are of interest in fMRI. Cases 1-2 show SAR close to 100% with fat suppression, which is reduced by a factor of 3 when fat suppression is avoided. Case 3 shows 30% better slice coverage (76 vs 60 slices) for a TR of 1500 ms. Cases 4-5 are examples of scans that can only be acquired without fat suppression. Note that SAR reduction depends on the pulses implemented in the sequence.

Skipping the fat-suppression pulse requires an alternative method to remove the lipid signal from the generated image. In this study, a reconstruction based on an SMS-like parallel acquisition reconstruction was demonstrated to separate water and lipid images. An extended formulation was outlined in this work to combine in a single formulation lipid-water separation, in-plane acceleration and SMS acceleration.

Lipid-water separation was compared to R_sms_=2 in Figure 2. The average g-factor only slightly increased (5%) for the lipid-water case, however, the standard deviation increased by x1.5. This rise in the standard deviation is because the main change in the g-factor maps is localized to the region of the shifted lipid. The SNR, as with SMS using CAIPIRINHA, is reduced when the g-factor is increased. In addition, the SNR of the acquired image actually improves when the fat-suppression pulse is removed, therefore one can further benefit from using this reconstruction method. Both phantom and human imaging showed an SNR improvement of ~30%. tSNR was estimated in a resting-state fMRI scenario and demonstrated a 12-50% increase upon removal of the fat-suppression pulse.

Total acceleration factors (*R*_*tot*_) of 2-16, including a x2 factor of the lipid-water separation, were demonstrated in phantom and in human imaging. Phantom imaging is commonly prone to larger artifacts due to its uniformity, yet, we were able to reconstruct high-quality lipid and water images. In human imaging, the lipid artifact – when using standard reconstruction without a fat-suppression pulse – can result in a local intensity deviation higher than 30%. The reconstruction method demonstrated here removed the lipid signal. However, combining R_PE_=2 and R_sms_=4 with lipid/water separation showed artifacts (Figure 5), which indicates potential limits of this method.

fMRI scans can benefit extensively from avoiding the fat suppression, which will reduce the SAR in 7T human imaging. In addition, the lower SAR can further be used to accelerate and shorten the repetition time or to include more slices. The current study demonstrates a reliable reconstruction of separate lipid and water images using the parallel imaging technique. However, it also introduces an additional factor that eventually competes for the limited resources available to accelerate the acquisition, defined by the sensitivity maps of the multi-channel coil. It can be seen in Figure 3 that the g-factor values are increased when the total acceleration factor is higher. Further research is required to optimize the FOV shift chosen for CAIPIRINHA when combined with lipid-water separation. In addition, one of the methods to reduce the parallel acquisition acceleration factor is combining the acquisition with the Compress Sensing^36^ method.

To further exploit the benefits of the lipid-water separation, lipid images can be utilized to improve movement detection. fMRI is sensitive to small movements, especially for long scans, and therefore detecting movements and correcting for them is vital. Thus, further research in our lab examines movement detection using the lipid image.

## Methods

### Phantom experiments

All scans in this study were performed on a 7T MRI system (MAGNETOM Terra, Siemens Healthcare, Erlangen, Germany) using a commercial 1Tx/32Rx head coil (Nova Medical, Wilmington, MA). The 3D phantom with brain and lipid compartments was 3D-printed based on Ref. ^37^. The inner compartment and the bottom outer compartment were filled with an agar mixture to mimic brain properties. The upper outer compartment was filled with peanut oil to mimic the precranial lipid layer. During the preparation, the phantom was gently rolled to generate a thin layer on the internal walls of the container to mimic the skin/muscle layer in the outer space adjacent to the lipid tissue. This phantom was designed to provide an RF field and B_0_ distribution similar to *in-vivo*, which is essential for 7T MRI tests. The agar mixture comprised 2.5% agar, 5.5g/L NaCl and 0.1mM GdDTPA.

Scans with the following parameters were performed to examine the reconstruction quality and to compare the images with fat suppression to those without it: FOV=220×220 mm^2^, resolution = 1.7×1.7 mm^2^ (130×130 pixels), slice thickness = 2mm, TR=2000 ms and minimal TE for each acceleration factor. Acceleration factors *R*_*PE*_=2 and *R*_*sms*_=2 (which are commonly called in Siemens software *R*_*PE*_ =2 and *R*_*sms*_=2) were examined. Two sets of gradient-echo (GRE) scans were collected to serve as input for sensitivity maps, one acquiring both fat and water (without fat suppression) and one obtaining the fat only, using water suppression. The common GRE scan parameters were FOV=220×220 mm^2^, in-plane resolution = 1.7×1.7 mm^2^, slice thickness = 2mm, TE=3ms. In the fat+water GRE, the scan specific parameters were: TR= 321 ms and total scan duration=0:42 min. In the fat only GRE, the scan specific parameters were: TR= 1790 ms, scan duration= 3:52 min.

To estimate the g-factor, based on the pseudo-replica method^38^, noise was randomly generated and added to the input signal 300 times. The g-factor was then estimated as the standard deviation per-pixel of the reconstructed images. In addition, to estimate SNR the scan was repeated 50 times and for each scan the signal was averaged over a central region (orange overlay in Figure 2). The SNR was then estimated from these 50 values by dividing their average by their standard deviation.

### Human scanning

Four human volunteers were scanned to examine the lipid-water separation in-vivo and to verify the fMRI efficiency. All methods were carried out in accordance with Weizmann Institute of science guidelines and regulations. All scans were performed according to procedures approved by the Internal Review Board of the Wolfson Medical Center (Holon, Israel) after obtaining informed suitable written consents.

The common scan parameters in Figure 4 were: FOV=208×208 mm^2^, slice thickness = 2 mm, TR =2000 ms, 30 slices, TE = 22 ms and 18 ms in set (a) and (b), respectively. Two GRE scans, one without fat suppression and one with water suppression were collected to serve as the input for generating the sensitivity maps. The common GRE scan parameters were FOV=208×208 mm^2^, resolution = 1.6 mm iso, TE= 3 ms. The fat+water GRE scan specific parameters were: TR= 273 ms, total scan duration=0:35 min. While in the fat only GRE scan, the scan specific parameters were: TR= 509 ms, scan duration= 1:06 min.

Figure 5 summarizes the experiment combining both in-plane and slice acceleration - R_PE_=2 with either R_sms_=3 or R_sms_=4. The common scan parameters for both sets were: FOV=220×220 mm^2^, 1.7 mm isotropic resolution, TE=28 ms, 60 slices. The TRs were TR = 1500 ms and 1000 ms for R_sms_=3 or R_sms_=4, respectively. The same GRE scans as in the previous paragraph were performed to generate the sensitivity maps. In addition, scan parameters for fMRI experiments were examined to identify SAR restrictive cases.

Figure 6 shows images acquired in the resting-state fMRI experiment. The SNR without fat suppression relative to the fat suppressed image was estimated at two representative regions (see orange overlays in Figure 5). It was calculated as the average signal within each region divided by the standard deviation in a noise-only region (outside the brain). The tSNR maps were calculated as the average (per-pixel) of the repeated scans divided by their standard deviation (per-pixel). The fMRI scan parameters were: FOV=220×220 mm^2^, resolution = 1.7×1.7 mm^2^, slice thickness = 1.7 mm, TR/TE = 1500/22 ms. The SAR level of the fMRI scans was examined with and without fat suppression.

## Acknowledgments

We are grateful to the Weizmann Institute of Science’s MRI technician team for assistance in the human imaging scans. Dr. E. Furman-Haran holds the Calin and Elaine Rovinescu Research Fellow Chair for Brain Research.

## Author contributions

Theoretical analysis and experiments were carried out by A.S. and R.S. E.F-H. and I.G. contributed in collecting the data.

## Additional information

Supporting Information is available in the online version of the paper.

## Competing financial interests

A.S. is employed by Siemens Healthcare Ltd, Israel, all other authors declare no competing financial interests.

## Supporting Information

### Extended parallel imaging formulation – including in-plane acceleration, inter-slice acceleration and lipid/water separation

#### 1.1 Two slices – No CAIPIRINHA

Let us assume we have two images, *I*_*1*_ and *I*_*2*_, of two different slices. Each image can be divided into a lower half *I*_*id*_ and an upper half *I*_*iu*_ (i = 1,2):

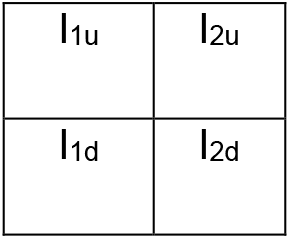

We acquire both images simultaneously, using a multiband excitation. Therefore, the measured signal is equivalent to:

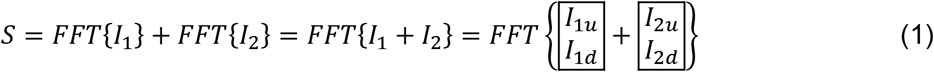

Now, assume we want to treat the problem as a parallel imaging problem, as if our final “full-FOV” image is:

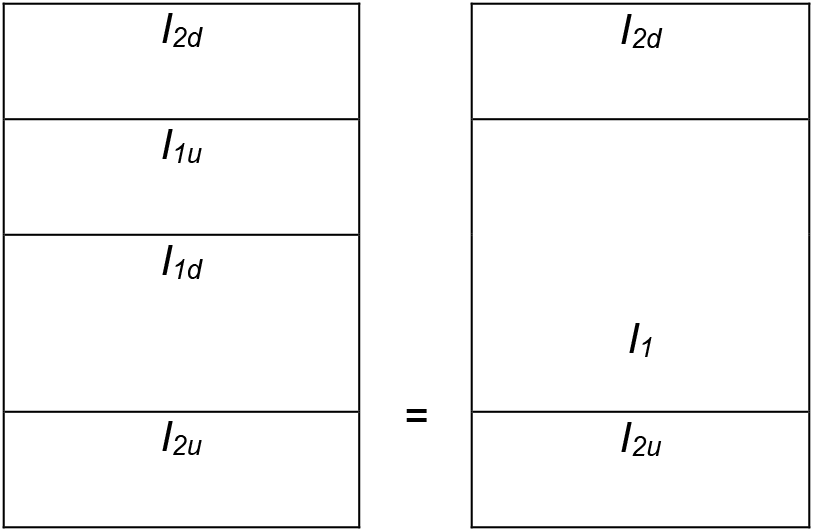

Note that *I*_*2d*_ (bottom half of image *I*_*2*_) is at the top, while *I*_*2u*_ is at the bottom. In this way, if we subsample (by a factor of 2) the FFT of this “full-FOV” image, parts *I*_*2d*_ and *I*_*2u*_ will wrap around to give the same image as in the SMS case.

To solve this new “parallel imaging” problem we need sensitivity maps of our “channels”. The sensitivity maps must match the “full-FOV” image (as in SENSE) and so must be of the form

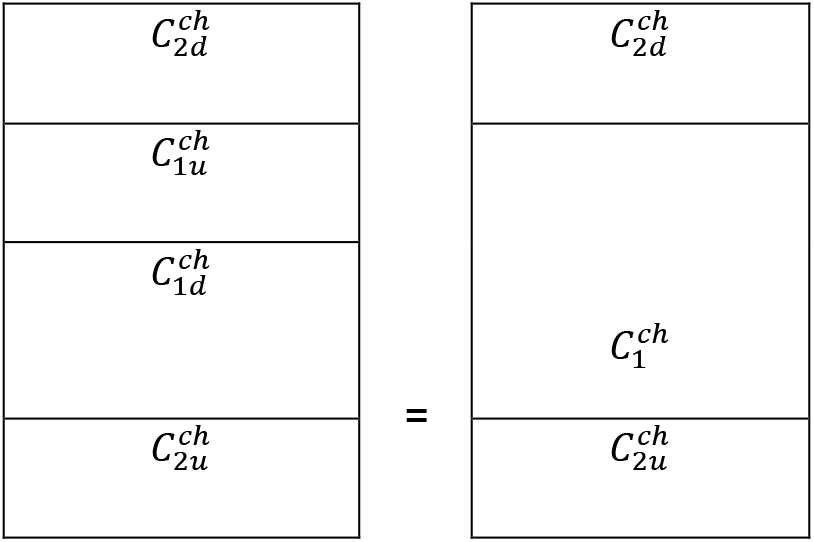

where 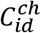 and 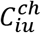 are the two halves of the sensitivity maps of channel *ch* and slice *i* (*i* = 1,2).

Finally, the signal can be described:

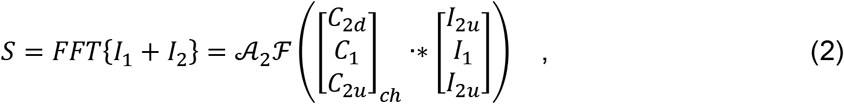

where 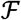 is an FFT operator and 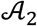 is a factor 2 subsampling operator that mimics the actual acquired dataset. After solving for the “full-FOV” image, the image can be split into the different slices.

#### 1.2 Three slices – No CAIPRINHA

For three slices we use the same principles as above, but we generate the “full-FOV” image as follows:

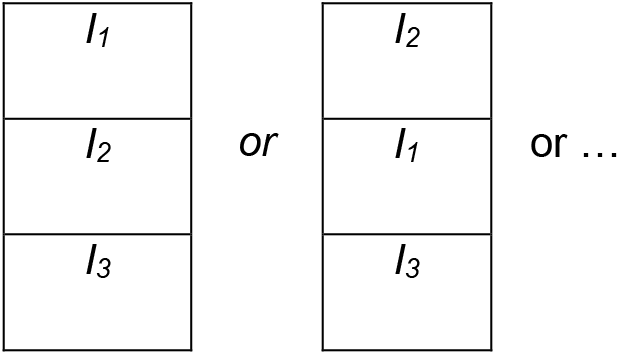

so that when sub-sampling by a factor of three the “wrapped” image will be the sum of all the slices. Note that here the images were not split into upper and lower halves, but rather used completely, since they wrap around completely. As a consequence, the effective sensitivity maps will be of the form:

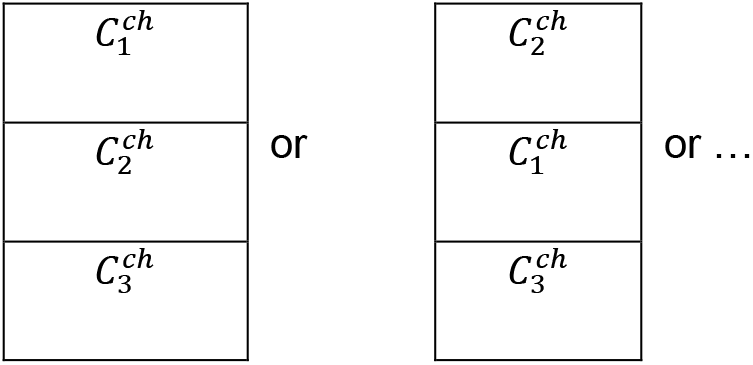

where 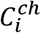 i is the sensitivity map of channel *ch* at slice *i* (*i* = 1,2,3).

#### 1.3 General case - No CAIPIRINHA

In the general case, we always create a “full-FOV” image that has one slice at its center (otherwise, some linear phase has to be added). In the odd case, that is simple; but in the even case, we always have to split one slice in half, putting one half at the top, the other at the bottom. Here are examples for four- and five-slice cases, with *I*_*1*_ always at the center, but the order can be arbitrary.

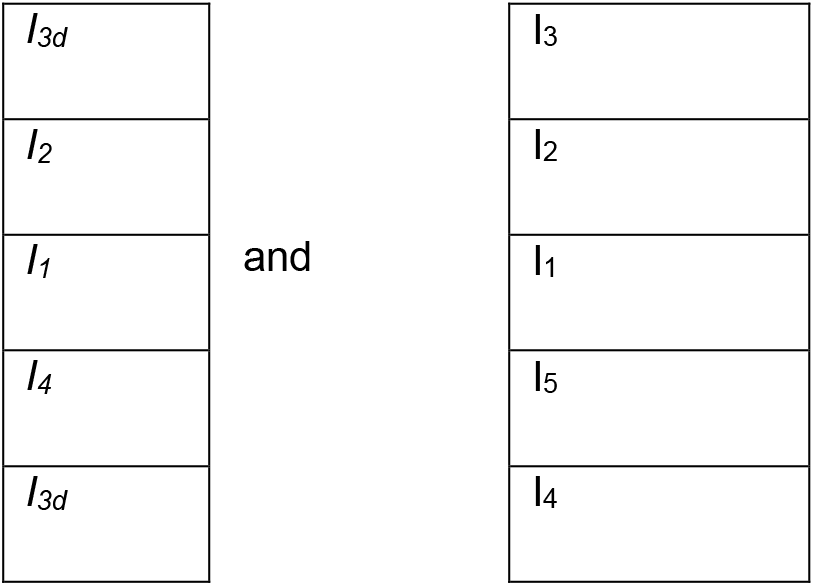

The channel sensitivities are analogous.

#### 1.4 CAIPIRNHA case

When using CAIPIRINHA, the same principles as above apply, but the effect of CAIPIRINHA on each slice must be taken into account: The image and sensitivity map within each slice undergoes a wraparound which gradually varies with the slice.

## Notes

### Competing Interest Statement

The authors have declared no competing interest.

